# DeepBend: An Interpretable Model of DNA Bendability

**DOI:** 10.1101/2022.07.06.499067

**Authors:** Samin Rahman Khan, Sadman Sakib, M. Sohel Rahman, Md. Abul Hassan Samee

## Abstract

The bendability of genomic DNA impacts chromatin packaging and protein-DNA binding. However, beyond a handful of known sequence motifs, such as certain dinucleotides and poly(A)/poly(T) sequences, we do not have a comprehensive understanding of the motifs influencing DNA bendability. Recent high-throughput technologies like Loop-Seq offer an opportunity to address this gap but the lack of accurate and interpretable machine learning models still poses a significant challenge. Here we introduce DeepBend, a convolutional neural network model built as a visible neural network where we designed the convolutions to directly capture the motifs underlying DNA bendability and how their periodic occurrences or relative arrangements modulate bendability. Through extensive benchmarking on Loop-Seq data, we show that DeepBend consistently performs on par with alternative machine learning and deep learning models while giving an extra edge through mechanistic interpretations. Besides confirming the known motifs of DNA bendability, DeepBend also revealed several novel motifs and showed how the spatial patterns of motif occurrences influence bendability. DeepBend’s genome-wide prediction of bendability further showed how bendability is linked to chromatin conformation and revealed the motifs controlling bendability of topologically associated domains and their boundaries.

## Introduction

Bendability is a critical mechanical property of genomic DNA with implications for its structure^1^, chromosomal packaging^2^, and interactions with DNA-binding molecules^3^. However, sequence signatures underlying DNA bendability are poorly understood at best. Advances in machine learning and Loop-Seq^4^, a SELEX-based assay of DNA bendability, offer an opportunity to address this gap. The recent Loop-Seq dataset, for example, quantified bendability across the entire yeast genome at 50 bps resolution (nearly 200,000 sequences). Loop-Seq starts with an initial library of PCR-amplified sequences. Each sequence has two flanking single-stranded overhangs that should attach to each other and form a loop if the sequence is naturally bendable. After leaving the library in a chemical solution for a specified time, the unlooped sequences are digested and the looped sequences are amplified for a second time. The experiment is then repeated using an identical library but omitting the digestion step. Finally, the bendability of each sequence is quantified as the logarithm of the ratio of its relative abundance in the two libraries^4^.

The first models of Loop-Seq data have been built from handcrafted features, such as, the frequencies and periodicities of dinucleotides^5^ and AT- or GC-rich sequences up to 6 bps in length^6^. These features are based on DNA biophysics and are easy to interpret, but the models leave ~60% of the variance in data unexplained. Furthermore, it remains unclear whether sequence patterns beyond these conventionally known dinucleotides and AT-/GC-tracts are important for DNA bendability.

To improve the state-of-the-art models’ performance in predicting bendability and discover the relevant sequence features *de novo*, here we introduce a convolutional neural network (CNN), DeepBend. Importantly, CNNs are known to suffer from a lack of interpretability^7^. To alleviate this issue, we designed DeepBend as a *visible neural network*^8^ with mechanistically grounded kernels that directly reveal: (a) the sequence patterns, also known as *motifs*, underlying DNA bendability and (b) how periodic occurrences or relative arrangements of motifs influence bendability.

This principle of designing individual components of a model to reflect domain knowledge has been recently discussed as *model-based interpretation*^9^. In general, model-based interpretation comes at the expense of lower performance than alternative “black-box” models^9^. Thus, we extensively benchmarked DeepBend on Loop-Seq data against alternative machine learning models, such as, support vector machines and random forests, and deep neural network architectures, such as CNNs and their hybrids with recurrent neural networks (RNNs). DeepBend consistently showed superior performance while retaining an extra edge in interpretability. Applying DeepBend on Loop-Seq datasets revealed both known and novel motifs and their relative arrangements important for bendability. The model also revealed the 7-mer GAAGAGC as a novel motif and its significant role in determining bendability beyond the conventionally known dinucleotides and AT-/GC-tracts. Finally, DeepBend revealed the sequence motifs influencing chromatin conformation through DNA bendability.

## Results

### DeepBend: A Deep Convolutional Neural Network (CNN) Model of DNA Bendability

DeepBend is a 3-layered CNN that takes in a one-hot encoded DNA sequence as input and predicts its bendability as output (Fig. 1). The first two layers are applied parallelly to both the forward and the reverse complement complement strand of the input sequence, allowing the model to detect patterns in both DNA strands. The first layer is a multinomial convolution layer with ReLU activation^10^. Filters in this layer detect motifs and the convolution operation computes matching scores of the motifs (log likelihood ratios, see Methods) at each position of the input sequence. The second convolutional layer learns spatial patterns in the motif matching scores. Up to this layer, all operations are applied to both the forward and reverse complement strands, thus producing two matrices. The element-wise maximum of the two outputs is passed to the next layer through a ReLU activation. This produces an output matrix which has matching scores of different spatial patterns of motif occurrences at different positions from both the forward and the reverse complement strands. In a post-processing step we identify the motifs with periodicity in their spatial patterns. The third layer is a single-filtered convolutional layer with linear activation. It takes in all the matching scores from the previous layer as input and outputs the predicted bendability of the sequence.

**Fig. 1.**
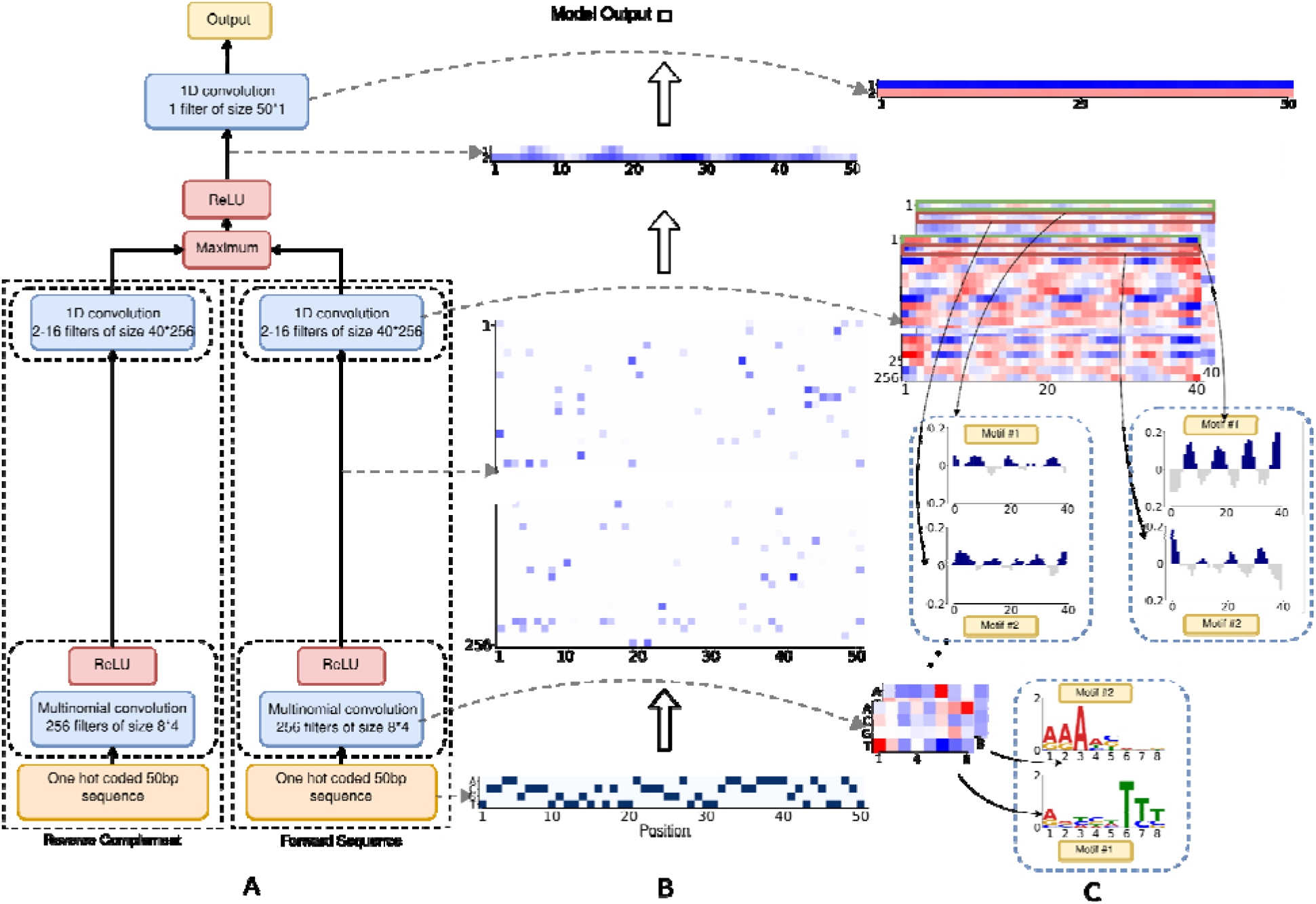
DeepBend model architecture and the inputs and outputs to different layers along with the convolution layers. **A-C.** One hot-encoded forward and reverse complement sequences are taken as input into the model. The first multinomial layer acts as a motif detector. After the first convolution and ReLU operation, we get a matrix containing the matching scores of the motifs at each position for both the forward and reverse complement sequence. The next convolution layer is designed to detect spatial patterns of motif occurrences and produces an output matrix for each sequence which contains the matching scores of these patterns at each position. The element-wise maximum of the results from the second layer are taken and then after a ReLU operation, we get as output the most prominent matches of bendability patterns at each position from both the sequences (forward and reverse complement). This is fed into the final convolution layer which gives the output bendability value. The first convolution layer provides motif patterns using transformation discussed in Methods Section 3, and their spatial patterns are obtained from the second convolution layer. The extraction of motifs and their patterns are shown in C.

Interpreting the DeepBend model is straight-forward. Since each row of a first layer filter is a multinomial distribution over the four nucleotides, these filters are directly interpretable as biophysical models of sequence motifs^10^. Regularising variance in the last layer separates out the relative spatial patterns of motifs significant for different ranges of bendability into different filters of the second convolution layer. These patterns can be easily identified from the weights of the second layer filters. Further details on the model architecture and the rationale thereof have been presented in the Methods section 2.

### DeepBend Models Loop-Seq Data with High Accuracy and Interpretability

We applied DeepBend to model four Loop-Seq libraries^4^, each quantifying the bendability of 50 bps long sequences: (i) The *Random library* has 12,472 randomly generated DNA sequences, (ii) the *ChrV library* comprises 82,404 sequences from yeast (*S. cerevisiae)* chromosome V (tiled across the chromosome with 7 bps shift), (iii) the *Nucleosomal library* contains 19,907 sequences centred at high nucleosome occupancy locations in the yeast genome, and (iv) the *Tiling library* consists of 82,368 nucleosome around 576 selected genes. Importantly, in the ChrV and Tiling libraries, each sequence overlaps with seven upstream and seven downstream sequences. For a fair and objective evaluation of the model performance, it is necessary to make sure that sequences in the test and the training sets do not overlap as this could otherwise inflate performance. Thus, while benchmarking on these two libraries, we have fit all models on specially designed datasets (please see Methods section for details). The training (test) sets obtained from this separation will be referred to as the *ChrV training (test) set* and *Tiling training (test) set*.

For comparing DeepBend with Basu et al.’s model^5^, we trained DeepBend on the Tiling library and tested it on the Random, Nucleosomal, and ChrV libraries. The Pearson’s correlation coefficient (r) between the true bendability of these three libraries and DeepBend’s predictions were 0.895, 0.931, and 0.774, respectively. Compared to Basu et al.’s models, the improvements were 59.82% in the Random Library, 55.16% in the Nucleosomal Library, and 29% in the ChrV library. The performance comparision between DeepBend and Basu et al.’s models^5^ have been shown in Fig. 2A. DeepBend also outperformed other machine learning models in terms of r: SVMs (by 118.29%), Random Forests (by 208.62%) and a series of deep neural network models (by 5.29-118.29%) (Supplementary Tables 1 and 2). All these models were trained on the Tiling library and tested on the Random library. We chose the Tiling library for training since it provides a comprehensive set of sequences spanning a whole chromosome, while the Random library was chosen for testing since the sequences are randomly generated and are equally likely to contain any 50 bps sequence.

**Fig. 2.**
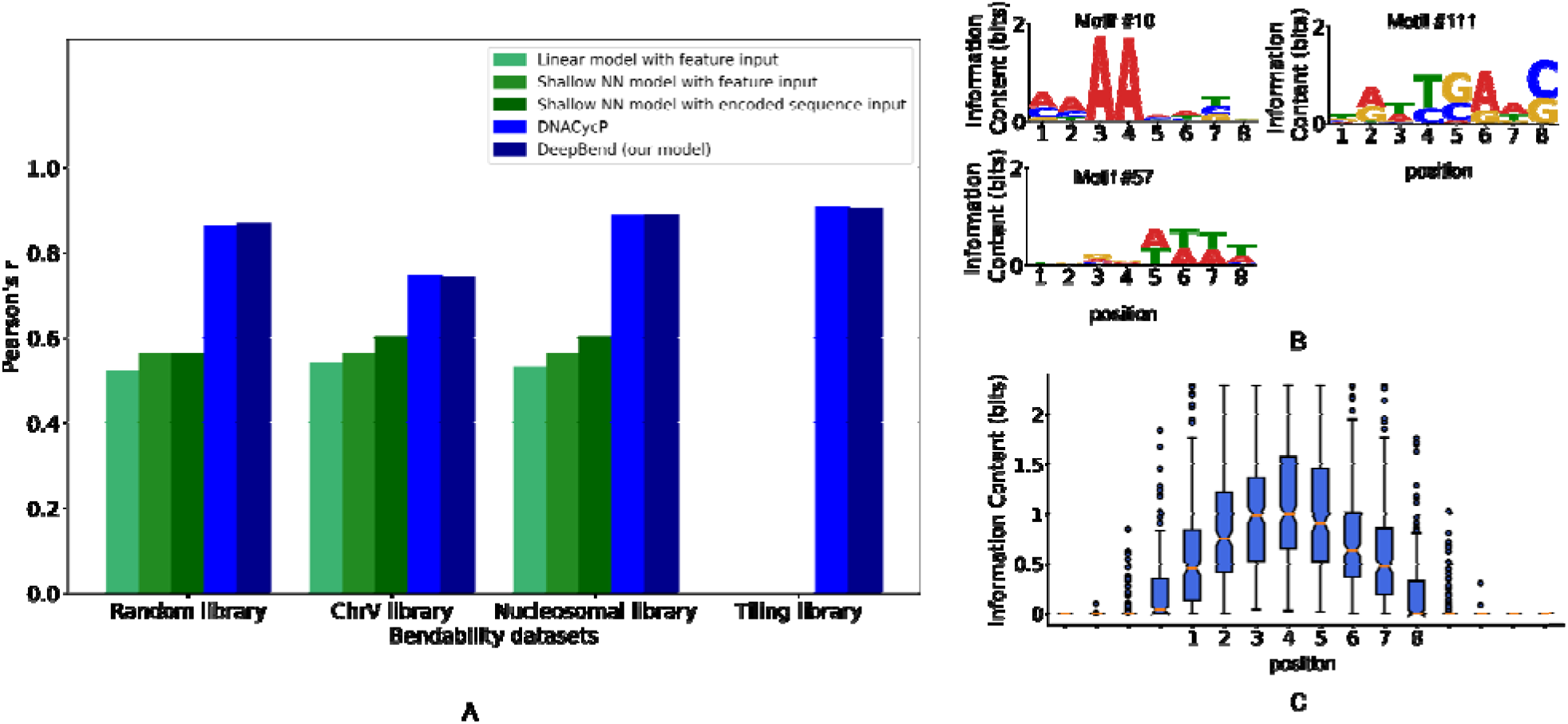
Comparison of model performance. **A.** Pearson’s correlation coefficient (r) between true and predicted results from different models. For DNACycP^11^ and DeepBend, 10-fold cross-validation results are shown for Random and Nucleosomal libraries. For ChrV and Tiling libraries, the results of the *ChrV test set* and *Tiling test set* are respectively shown. For the other three models, we have shown Basu et al.’s^5^ results of testing their model on the Random, ChrV and Nucleosomal libraries. Since they trained models on the Tiling library, we could not show their performance considering the Tiling library as test data. **Motifs from DeepBend. B.** Motifs 10, 57 and 111. Motif #10 and #57 are previously known bendability motifs and motif #111 is a novel motif. **Information content of motifs from DeepBend. C.** Distribution of information content of the motifs at each position, showing that motifs are small in size, of length 3-5. The motifs were centred.

During the preparation of this manuscript, we came across Li et al.’s DNACycP model^11^, which is a hybrid of CNNs and RNNs, for modelling Loop-Seq data. Although DeepBend outperformed RNN models in our benchmarking (Supplementary Table 2), we still performed a rigorous comparison between DeepBend and DNACycP as follows. For the Random and Nucleosomal libraries, we compared the models using 10-fold cross-validation. For the ChrV and Tiling library, we trained and tested the models on the specially designed datasets noted above ensuring that sequences in the training and the test sets do not overlap in the yeast genome. In all these comparisons, DNACycP and DeepBend showed nearly identical performance (Table 1). The performace comparison of DeepBend and DNACycP are also shown in Fig. 2A. The Pearson’s correlation coefficient (r) between the true bendability and DeepBend’s predictions for models trained and tested on different libraries are provided in Supplementary Table 3.

**Table 1.** Performance comparison between DeepBend and DNACycP. Shown are the Pearson’s correlation coefficient (r) between true and predicted values.

| Test Libraries | DeepBend | DNACycP |
| --- | --- | --- |
| Random Library<sup>A</sup> | 0.87 | 0.86* |
| Nucleosomal Library<sup>A</sup> | 0.89 | 0.89* |
| ChrV Test Set<sup>B</sup> | 0.74 | 0.74 |
| Tiling Test Set<sup>B</sup> | 0.90 | 0.91 |
| * Results from original paper<sup>11</sup>. <sup>A</sup> Results from testing during 10-fold cross-validation.<br><sup>B</sup> Results from testing on the specially designed test set. |  |  |

### DeepBend Confirms Known Motifs and Discovers New Motifs Influencing Bendability

Visualizing DeepBend’s first layer kernels we found that the model was able to learn the motifs that are conventionally known to influence bendability^5,6,12–14^. Interestingly, DeepBend also found some new motifs and predicted a significant role for them in determining bendability. We ranked DeepBend’s motifs according to their contribution to positive and negative bendability, as we quantified from the change in model’s prediction after deactivating one motif at a time (Methods 4). From all the DeepBend models we have trained, for interpretation purposes we have selected the model that has been trained on the ChrV library and has a reduced number of filters (two) in the second convolution layer. All the motifs from this model along with their patterns have been provided in Supplementary Material 1. Among the known motifs are the A/T dinucleotides (Motif #3, #10, #178, #232), A/T regions (Motif #8, 57, 150, 166, 213, 250), C/G regions (Motif #34, #24) which contribute to bendability positive or negatively depending on their relative arrangements ^5,6,12–14^ (discussed below). Although the visually discernable “core” signals from these motifs are similar, they differ in the flanking regions around the core signals. Likewise, Poly A tracts (Motif #11, #51, #117)^15^, CG (Motif #28) are negatively contributing to bendability in general. Among the novel motifs there are motifs like GAAGAGC (Motif #40) and Motif #47 which contribute positively, and motifs like Motif #78, Motif #111, Motif #77 and Motif #99 contribute negatively. The most prominent is motif #40, heptamer GAAGAGC, which is the most informative motif found by our model and is strongly indicative of positive bendability. Some examples are shown in Fig. 2B.

Expectedly, most of the specific motifs are small, usually dinucleotides, as the bendability of the sequence is fundamentally determined by the bendability between adjacent bases. From the distribution of information content at each position of the motifs after centring them, we find that the average information content is much less than the maximum value and that most motifs are small in size, of length 3-5 (Fig. 2C). So, in effect, DeepBend is suggesting that bendability of a sequence is principally determined by short motifs and arrangements thereof.

As has been shown in ^4^, nucleosome regions have a peak in bendability near the dyad (Fig. 3A). This definite pattern occurs due to the presence of bendability motifs around these regions. In nucleosomes, generally, A/T dinucleotides occur at a 10 bp period in antiphase with G/C dinucleotides. Most of the motifs identified by DeepBend are distributed at different distances from the centre of the nucleosome. These motif distributions around nucleosomes conform to prior knowledge ^16,17^. Motifs having dA:dT (motif #49, motif #50) are found less in the nucleosomal region. In addition to this, some more diverse motifs found by DeepBend are also specifically positioned around nucleosome regions, which can explain the bendability around nucleosomal regions further. For example negatively contribuitng Motif #245 is found less at the centre of the nucleosomal region, whereas there is sharp increase and decrease in the presence of positively contributing Motif #137 at the centre of nucleosomal regions. Examples are given in Fig. 3B. So, the motifs from DeepBend can be further used to explain bendability around nucleosomal regions.

**Fig. 3.**
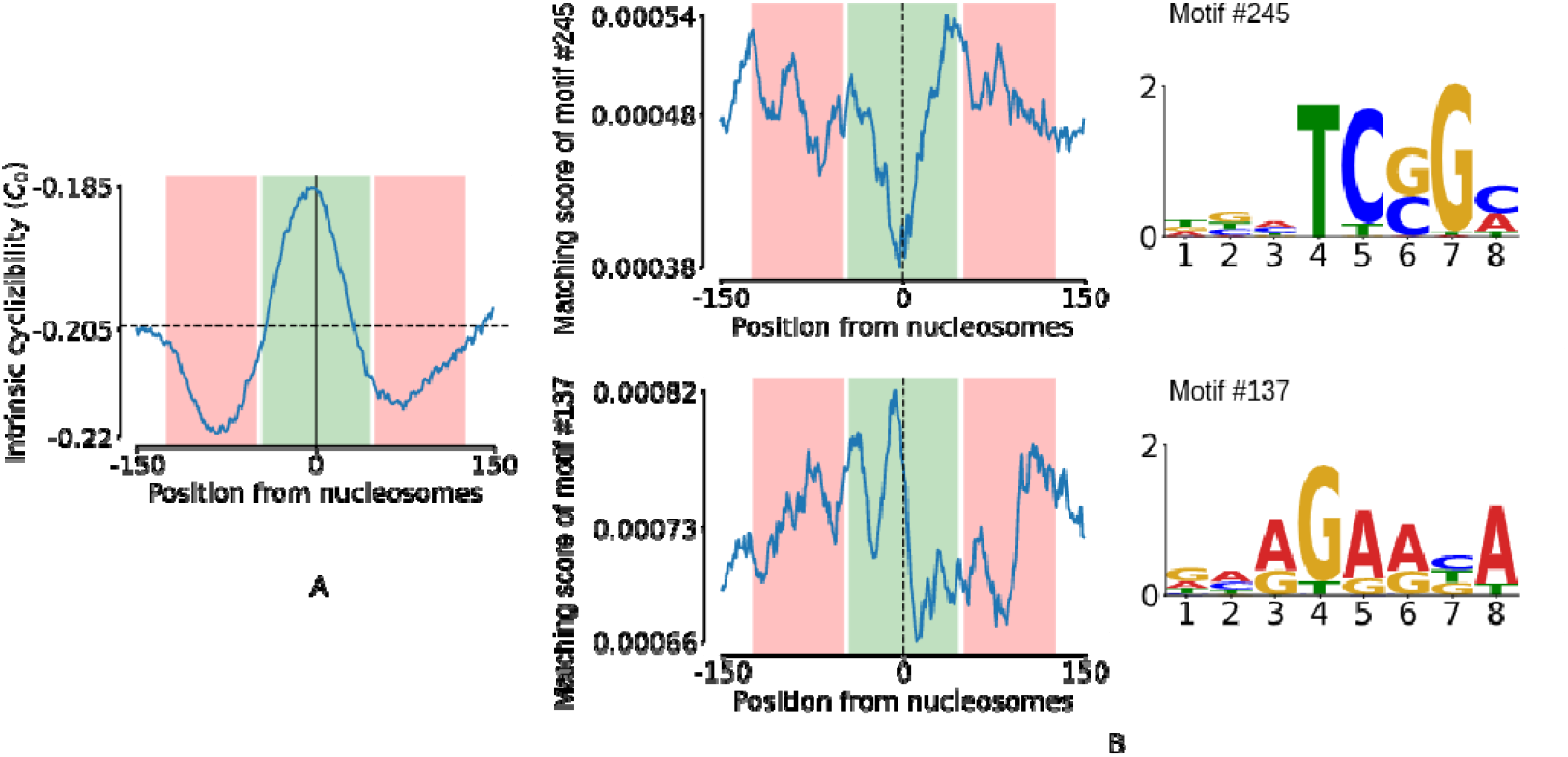
Presence of novel motifs around nucleosomal regions. **A-B.** A. Average bendability around nucleosomes (the green region from +45 --45 is more bendable compared to +-125 - +-50 region) and B. the strong/weak matching scores (Methods 5) of motif-245 show that it is found less in the central region, which is expected since motif #245 is a negative contributing motif, which can be observed from its spatial pattern and also by global importance analysis as discussed in Methods Section 6. Similarly, positive contributing motif #137 is found more abundantly in the central region.

### DeepBend reveals spatial and periodic patterns of motif occurrence

DeepBend can learn how the motifs act throughout the sequence without requiring any prior information. DeepBend captures important relative spatial patterns of small motifs in sequence segments rather than fixed features such as k-gapped dinucleotides or k-mer counts.

#### The second layer filters capture spatial biases or periodic patterns in motif occurrences

Each of the rows of the filter corresponds to the relative spatial pattern of a motif from the first layer. We call each such pattern a first-order pattern. The periodic presence of a motif is an example of a first-order pattern. Due to the property of convolution, we can know how the motifs should be distributed for a particular pattern to be present from the weights of the filters of this convolution layer. Fig. 4 presents a pictorial description of how DeepBend detects spatial patterns. Here, the filter row or kernel corresponding to a motif from the previous layer for positive bendability are shown. The regions where the motifs should be (relative to each other) have positive values and are shown in blue; the regions where the motifs should not be (relative to each other) have negative values and are shown in grey (Fig. 4A-C), and the magnitudes of the weights provide the significance of the presence or absence of the motif at that position.

If there is a spatial bias or periodicity between the occurrences of several first-layer filters, the filters in the second layer can also learn that. Fig. 4A shows that the periodic patterns of motifs #24 and #192 are interleaved with one another with an offset. Such an interleaving periodic presence of the two motifs greatly increases the bendability of the region. We refer to such composites of first-order patterns as higher-order patterns. Each of the filters from the second layer is actually a higher-order pattern. Thus, the number of filters in the second layer determines the number of different higher-order patterns the model can learn. By hyperparameter tuning, we have noted that two filters in the second layer are sufficient. This is because the first-order patterns of high and low bendability can co-exist separately in their respective single filter. We have seen only two main types of first-order relations: (1) in-phase and out-of-phase periodic patterns and (2) patterns that remain consistently high or low throughout a sequence. To make each of these filters in the second layer capture patterns that are representative of a bendability throughout the sequence, we have added position-wide variance regularisation in the third layer corresponding to each filter in the previous layer. This allows us to easily separate out the patterns for high and low bendability.

Instead of capturing long motif patterns, the model is able to learn spatial patterns through which it can find larger patterns with smaller motifs. Because of this, it is more expressive than models that take fixed gapped motifs as features. The periodic patterned distributions from the second layer, seen more often for positive bendability patterns, are useful for finding the presence of gapped motifs. An example is shown in Fig. 4A. The pair of motifs #24 and #190 occurs at a period of ~10bp at the centre of the example sequence. These periodic patterns are captured by the 2nd layer convolution when the window passes over this region, producing a high output there.

The motifs and their first-order patterns learnt by our model can be summarised in a table like that shown in Fig. 4B,C. This model has been trained on the ChrV Library and has 2 filters in the second convolutional layer. The motifs and their relative arrangements that are important for positive and negative bendability are shown here. The motifs on the left are some confirmatory motifs that show that from motif #3, the model learns that periodic occurrence of TT dinucleotide at a helical distance of 10bp can make a sequence more bendable, which was also reported by ^5^. Periodicity of T/A dinucleotides at ~10bp is also considered as a preferred sequence for nucleosome positioning ^16^ and this high bendability explains how these sequences loop around nucleosomes. Motif #232 from our model tells us that the presence of a short sequence of alternating Ts and As makes sequences more bendable regardless of their distribution. ^5^ also reported TA dinucleotide to have the highest correlation with positive bendability among all dinucleotides. Periodic presence of A/T regions and C/G regions makes sequences more bendable ^5,6,14^. The first-order periodic patterns for high bendability for motif #0, #8, #23, #34, #150, #183, #229 from our model also show this. Observing the higher-order relations between these regions shows that an interleaving periodic presence of these A/T and C/G regions makes the sequence even more bendable. Interestingly, the same motif can contribute both positively and negatively depending on how they are arranged. For example, Motif #196 contributes positively when periodically arranged but contributes negatively when arranged more consistently.

In addition to the confirmatory motifs discussed above, DeepBend has also found some novel motifs. The distribution of motif #43 reveals that CTGG sequence is positively contributing when positioned at helical distances. In general, positively influencing motifs mostly act at periodicities of the helical length of DNA and are small and specific. Negatively influencing motifs do not show these periodic patterns and are mostly non-specific/diverse. All the motifs from our model and their patterns are presented in Supplementary Materials 1.

Higher-order relative patterns can be visualised by overlaying the first-order patterns of multiple motifs over each other (Fig. 4C). We found the relative tendency of two motif distributions to activate together for high bendability and observed the higher-order patterns for the highest-ranked motif pairs. As expected, similar-looking motifs activate together the most and their patterns overlap completely. Different-looking motifs also activate together for high bendability and provide more interesting information. The higher-order patterns of such a motif pair reveal how these motifs are arranged relative to one another. The second example in Fig. 4C shows that periodic A/T regions interleaved with periodic G/C regions in anti-phase increase the bendability. So we see that from the second layer of DeepBend, it is possible to easily determine both signifcant first-order and high-order relative arrangement of motifs.

### DeepBend Revealed the Novel GAAGAGC Motif and Its Strong Role in Determining Bendability

As noted above, DeepBend has learned several novel motifs beyond the conventionally known poly-A or dA:dT patterns. Of these, the GAAGAGC motif (motif #40) (Fig. 4C) is particularly interesting. The 7-mer GAAGAGC is present in many viral genomes, sometimes highly conserved within a hyper-variable region and potentially has a role in RNA folding and viral pathogenicity^18,19^. However, this motif’s connection with DNA mechanical properties like bendabilty has never been discussed.

To substantiate the GAAGAGC motif’s role irrespective of our model, we checked the *bendability quotient* of the 7-mer GAAGAGC and its reverse complement, GCTCTTC. Basu et al. defined and used this statistic to denote the frequency of a k-mer’s occurrence in sequences with high bendability compared to those with low bendability^5^ (Methods). Importantly, GAAGAGC and GCTCTTC showed the highest bendability quotient of all 7-mers (Fig. 5A), indicating a clear role for this motif in making the DNA more bendable. Furthermore, GAAGAGC is much more bendable than the 7-mers that differ with it by even a single nucleotide. The bendability values of ChrV sequences containing GAAGAGC are significantly more positive than those containing 7-mers mismatching GAAGAGC (Fig. 5B). Besides, GAAGAGC and GCTCTTC occur 120 times in ChrV, which makes them among the top 14% 7-mers found in ChrV (Fig. 5C).

**Fig. 4.**
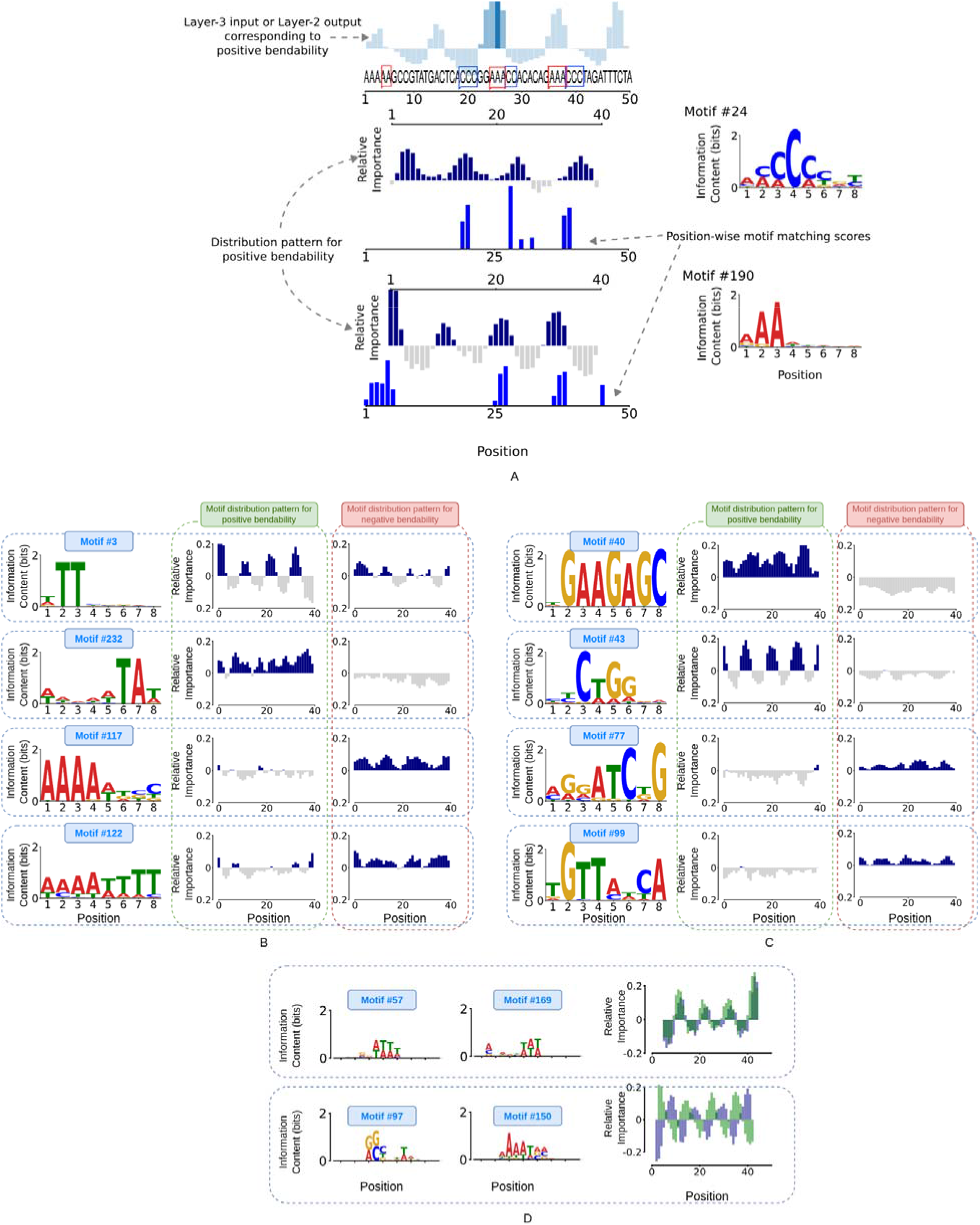
Model detecting spatial patterns. **A.** ChrV:33804-33853 is a highly bendable sequence. The input corresponding to the row with positive weights in the third layer shows the window positions where the second convolution layer has detected patterns of high bendability. When the convolution overlaps at a position where there is a good match between the spatial patterns and the matching scores of the motifs, as shown in the figure there is a high output from the second layer at that position for that filter. The position highlighted in darker blue shows a high output due to the presence of positively contributing distributions of motifs in this region at close proximity. Motif #24 and motif #190 are distributed periodically, which is an indicator of positive bendability matching the first-order patterns of the filter. And also these distributions are interleaved at the same region making that region more bendable, matching the higher-order relative arrangement of the motifs as well, which further intensifies the signal for positive bendability. **Shows some motifs found by our model and their relative distribution for positive and negative bendability. B-C.** B. DeepBend has confirmed that simple motifs like TT(#3) and TA(#232) contribute positively when they are at periodic helical length distances of 10 bases. Periodic TAs (#232) also contribute positively throughout the sequence. Long A(#117), T and dA:dT(#122) contributes negatively. C. In addition to more traditional motifs, DeepBend is able to capture more elaborate motifs like #40, #43, #77, and #99. **Higher-order relation between first-order spatial patterns of motifs. D.** Each row shows a pair of motifs that have been centred and their overlayed spatial patterns for high bendability after correcting for the offset. Similar looking motifs activate similarly and their patterns also overlap. DIfferent motifs like #97 and #150 also activates in unison for high bendability. These motif distributions, if placed at ~10bp periodicity with a 4-5bp phase difference from each other, are most bendable.

Given such frequent occurrence of the GAAGAGC motif and its potentially strong connection to bendability, we asked whether this motif plays a role in biologically important regions, such as, around Transcription Start Sites (TSS). Basu et al first showed the presence of a sharp rigid region upstream of TSS^4^. We determined mean bendability of −600bp to +400bp regions around TSSs in all 16 yeast chromosomes along with the occurrence pattern of GAAGAGC/GCTCTTC. We found that the occurrence of GAAGAGC/GCTCTTC drops in the rigid region and peaks in the bendable region, supporting its influence on bendability around TSS, Fig. 5D.

**Fig. 5.**
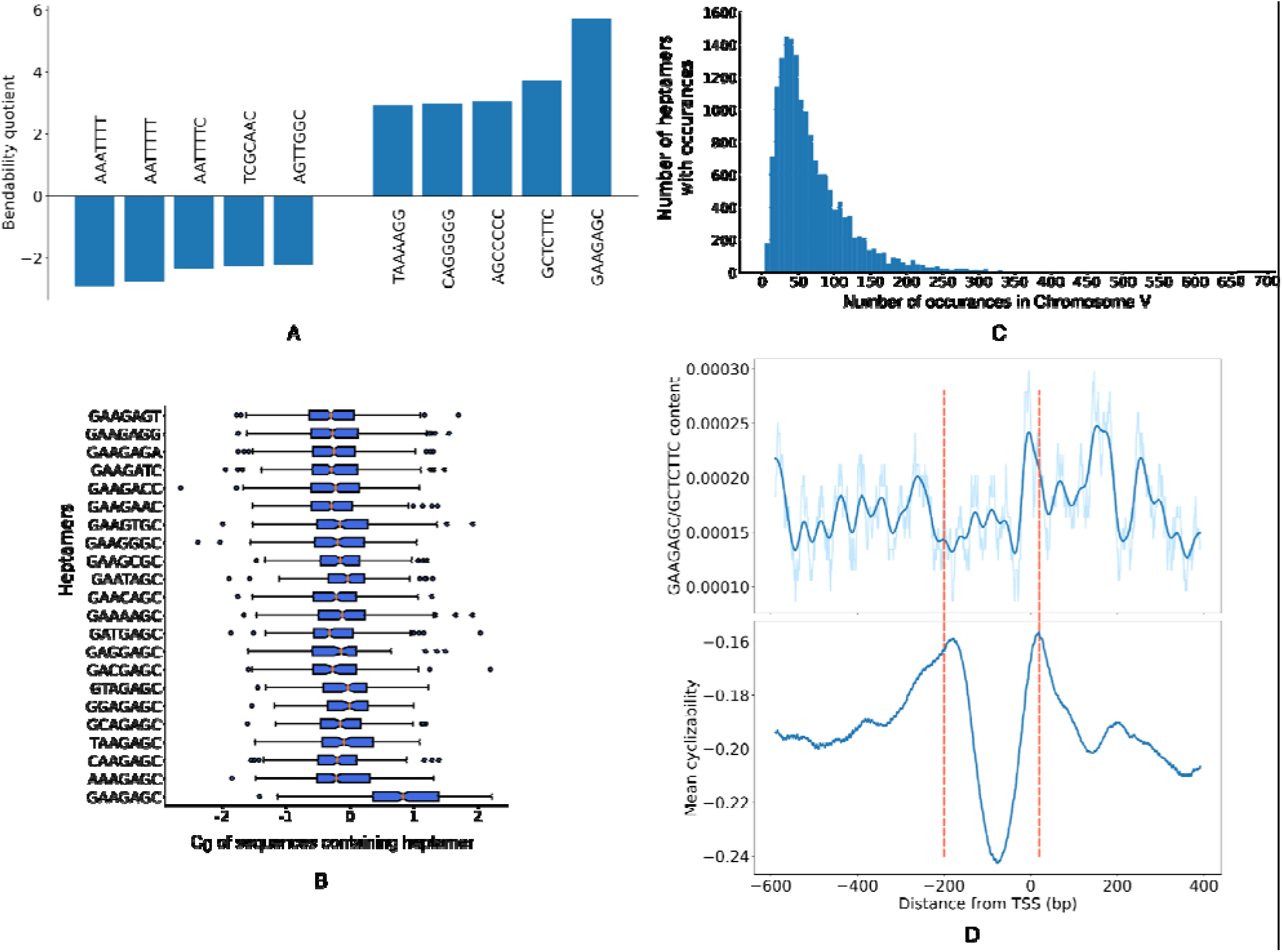
GAAGAGC Motif. **A-D.** A. Bendability quotient of heptamers with highest 5 and lowest 5 values. B. Boxplot showing the distribution of the bendability scores of the sequences from ChrV library containing GAAGAGC and its neighbours (which differ from GAAGAGC by only 1bp). C. Shows the distribution of heptamers occurring at different counts in ChrV. D. GAAGAGC/GCTCTTC content and predicted mean bendability around (−600bp-400bp) all TSS in 16 chromosomes of *S. Cerevisiae*.

### DeepBend Reveals Bendability Property and Sequence Motifs Associated with Chromatin Conformation

Eukaryotic chromosomes are organized into Topologically Associated Domains (TADs) where sequences within a TAD are more likely to interact with each other compared to sequences across TADs. As we reviewed below, recent studies found several molecular covariateds of TAD formation in yeast. However, it has not been reported whether specific sequence motifs influence TAD formation in yeast through controlling DNA bendability. CTCF binding sites are found to form boundaries between TADs in metazoans, and CTCF binding sites were found to have higher DNA bendability than surrounding DNA in five different species^11^. Notably, CTCF is not present in yeast, and the location of TAD in yeast has been explained with nucleosome positioning and histone marks^20^. Nucleosome-resolution mapping of chromosome folding revealed that boundaries separating domains in yeast are strongly enriched for the nucleosome depleted regions (NDRs) or long linkers (the sequences between consecutive nucleosomes) that are often found in yeast promoters^20,21^. However, not all NDRs form boundaries and strong boundaries tend to occur at promoters of highly transcribed genes. Some other features of yeast boundaries include enrichment of a variety of histone marks such as H3K4me3 and H3K18ac, and high levels of the RSC ATP-dependent chromatin remodelling complex and high levels of the cohesin loading factor Scc2^20^.

To determine if bendability plays a role in TAD formation and identify the motifs that influence TAD formation, we predicted the bendability of all 16 yeast chromosomes using DeepBend and analyzed a Hi-C matrix of yeast chromosomal contacts at 200bp resolution^22^ (Methods 7). Our analysis revealed a clear connection between boundary formation and bendability (Methods 6). Comparing the bendability of ±500 bp regions at boundaries against ±500 bp regions at domain sections (Methods 8), we found that boundaries have a 120 bp rigid region at the center flanked by two ~400 bp highly bendable regions (Fig. 6A,B). To further corroborate this finding, we sorted the boundaries into quartiles of their TAD separation score^23^ and analyzed DeepBend’s predicted bendability values of ±200 bp at the boundaries. Indeed, as we consider boundaries with weaker separation scores, bendability increases at central regions and while the linkers become shorter (Fig. 6D,E). Finally, promoters at boundaries are more bendable on both sides compared to promoters at domains (Fig. 6F).

**Fig. 6.**
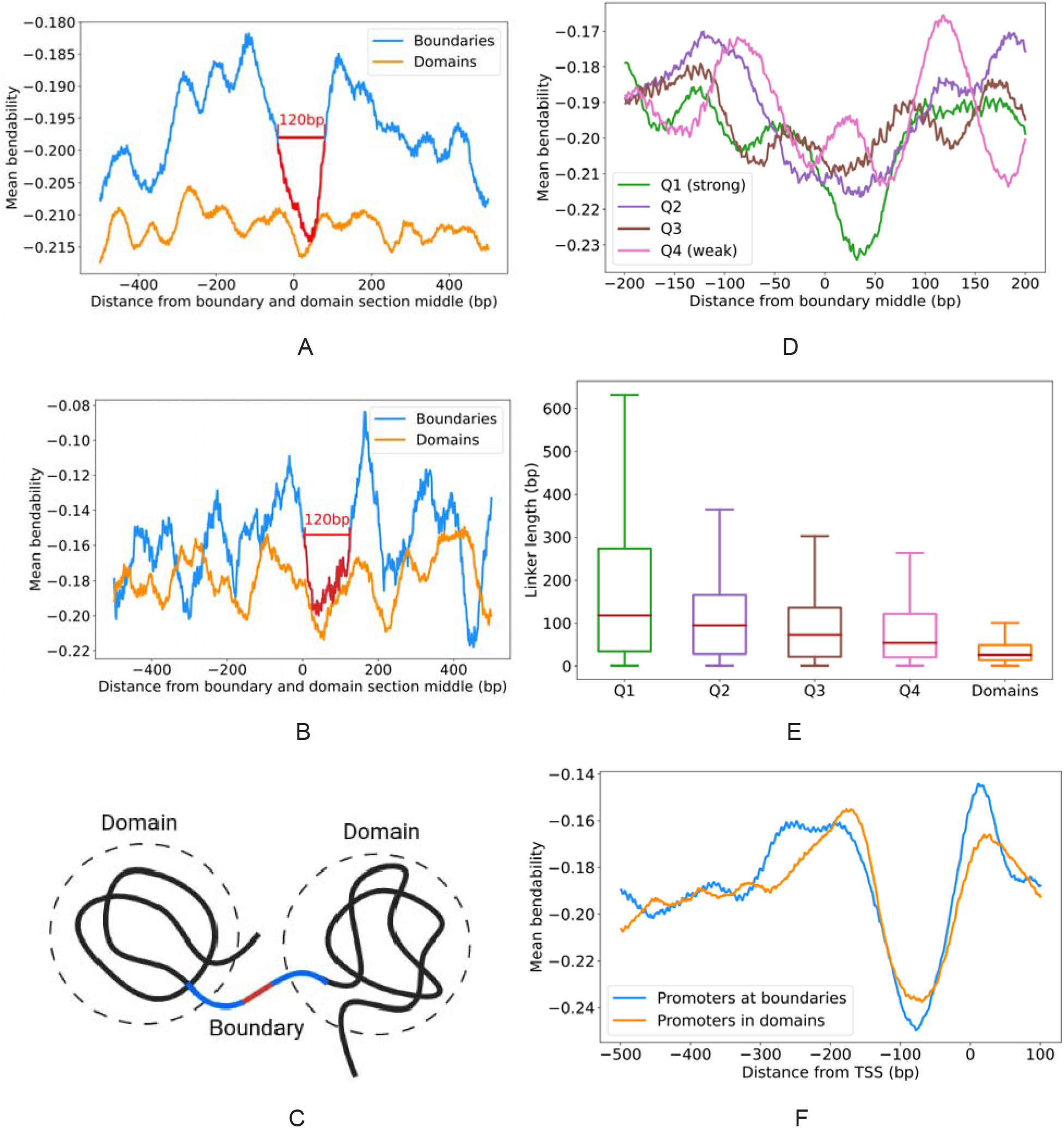
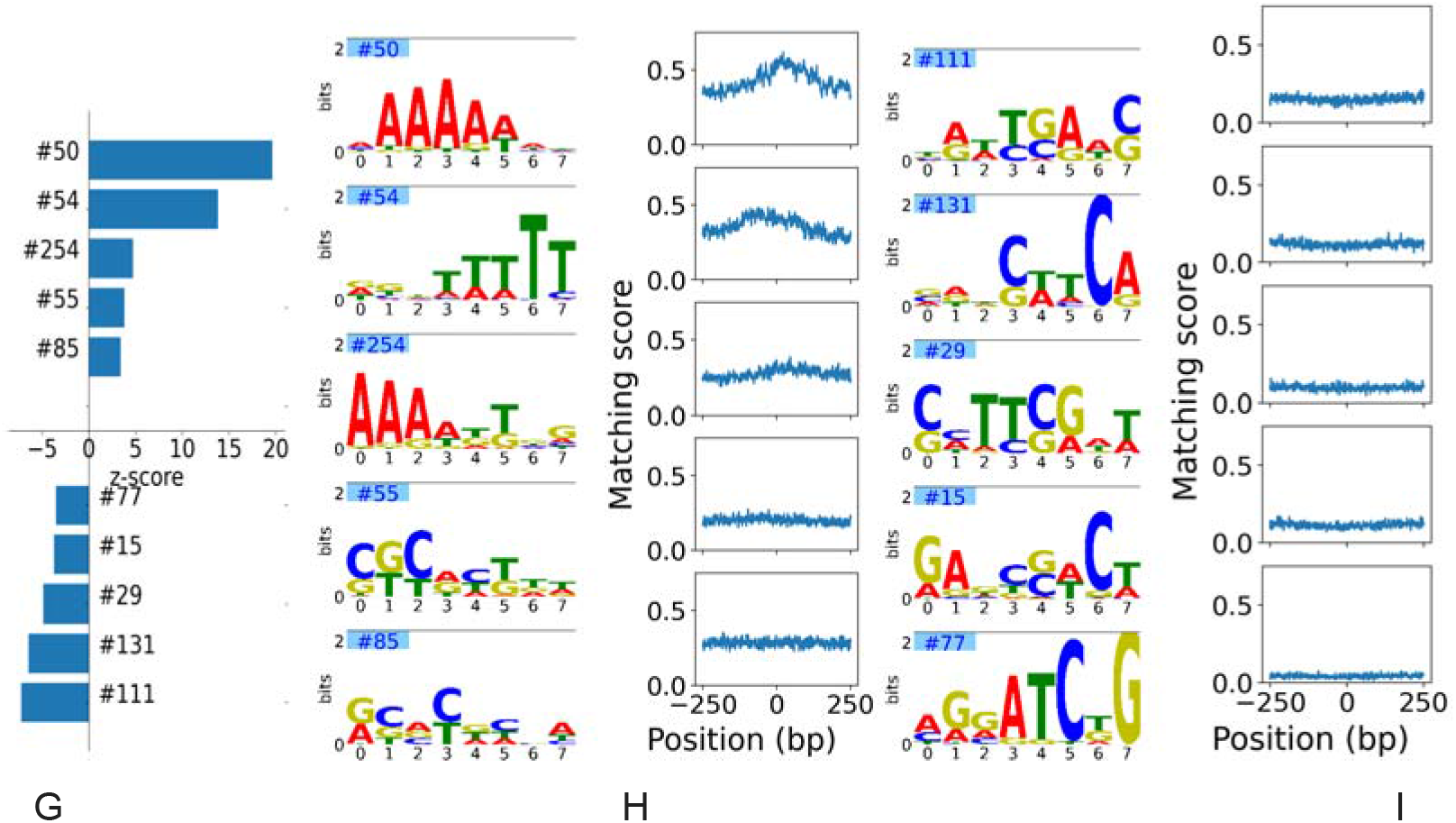
Stronger boundaries are more rigid. A. Mean bendability of 1000 bp regions at boundaries and 1000 bp sections of self-interacting domains of all 16 chromosomes where bendability is predicted by DeepBend. B. Mean bendability of 1000 bp regions at boundaries and 1000 bp sections of self-interacting domains in chromosome V where bendability data is obtained from Chromosome V library. C. Formation of boundaries in yeast. A stiff region (red) forms the main part of the boundaries which is flanked by two highly bendable regions (blue). D. Boundaries in all of the 16 chromosomes in yeast are split into quartiles (Q1-Q4) based on their TAD separation score. Then, bendability of all boundary regions in a quartile was averaged. E. Distribution of lengths of nearest linkers from boundary middle positions of quartile boundaries and rest of the linkers that are considered in domains. Boundaries from all 16 chromosomes were considered. F. Promoters at boundaries contain rigid NDRs flanked by highly bendable regions compared to promoters at domains. Regions between 500 bp upstream and 100 bp downstream of transcription start site were considered promoter regions. Bendability of promoter regions from all 16 chromosomes were averaged. A promoter was considered at a boundary if the middle of any boundary falls in this promoter. G. Z-score of some motifs that influence bendability more in boundaries compared to domains (top) and in domains compared to boundaries (bottom). H. Motif logos and mean matching score of 5 motifs with positive Z-score in G. I. Motif logos and mean matching score of 5 motifs with negative Z-score in G.

Next, we investigated which sequence motifs are relatively more important for determining bendability in domains and boundaries. We first calculated the motif matching score, a score derived from DeepBend denoting the relative match of the learned 256 motifs at some position of a DNA sequence, in all 16 chromosomes of yeast (methods 5). For each motif, we conducted a two-sample z-test^24^ for each motif to compare its presence in domains and boundaries in terms of the distribution of its matching scores. We considered matching scores in −250 bp to 250 bp around each boundary middle as distribution of motifs in boundaries and scores in the remaining chromosomal region as distribution in domains. A positive z-score indicates both higher presence and influence in determining bendability in boundaries compared to domains. Z-score also allowed us to create a ranking for all 256 learned motifs (Supplementary Tables 5). In general, sequence motifs containing poly (dA:dT) showed higher z-scores denoting that they highly influence bendability in boundaries (Fig. 6H). Besides these classes of sequences, motifs with lower, but still significant positive z-score values contain poly (dC:dG) sequences that are likely to influence bendability at boundaries, albeit to a lesser extent than poly (dA:dT) sequences. Motifs that have the lowest negative z-scores, denoting higher influence on bendability in domains, consist of shorter sequences of A/T and C/G, such as, TT and TC dinucleotides (Fig. 6I). Our finding conforms with the fact that homopolymeric sequences, poly (dA:dT) and poly (dG:dC) are prevalent in promoters of *S. cerevisiae* ^16^ where boundaries are more likely to occur and short A/T and G/C sequences are more likely to occur in periodic nucleosomal sequences which are less found in boundaries. Among the motifs with significant z-scores, we show some motifs in Fig. 6 that represent the sequence patterns that influence bendability in boundaries and domains. We also ran a motif matching tool, streme ^25^, to find out relatively enriched motifs in domains and boundaries. The motifs found by streme are similar to our motifs with significant z-scores (Supplementary Notes 2).

From the mean matching score patterns of these motifs in −250 bp to + 250 bp around boundaries, we found that prevalent motifs in boundaries, like those containing long A/T sequences, tend to occur more at the boundary centres and gradually less at positions farther from boundaries. Less prevalent motifs in boundaries appear less throughout the surrounding region around boundaries. Prevalence of poly (dA:dT) sequences might create the sharp rigidity in boundaries shown in figure 5A^16^.

## Discussion

DeepBend is able to match the predictive performance of the best models on large scale Loop-Seq datasets along with providing an unified explanation for this performance. Interpretability has spurred the development of post-hoc interpretation methods to infer the sequence motifs potentially driving CNN’s predictions. Such post-hoc methods are complicated to use. They involve choosing thresholds for several parameters but the right approach to do this is still unclear. Furthermore, the efficacy of their discovered motifs in modelling the data has not been systematically assessed. Finally, it is complicated to check higher-order patterns if they are spread over multiple layers in the model. DeepBend is able to provide the first-order and higher-order spatial patterns responsible for high and low bendability thanks to design choices like using a wider second convolution layer and adding positional variance regularisation in the final convolution layer.

DeepBend improves over previous interpretable feature-based models. Such improvement can be attributed to two aspects as follows. Firstly, DeepBend is capable of capturing continuous distributions - it captures the spatial patterns of small motifs in sequence segments rather than fixed features, such as, k-gapped dinucleotides or k-mer counts. Convolution captures the spatial relations better. Secondly, DeepBend is capable of identifying more complex probabilistic motifs that are not possible to be captured in simple models: a 3-layered CNN model like ours is able to learn more complex motifs that are not possible in simpler models.

The motifs found by DeepBend include both known and novel motifs. GAAGAGC and its reverse complement GCTCTTC are the most specific motifs found by our model and indeed these are the most bendable 7-mers. DeepBend reveals that stronger TAD boundaries tend to have more rigid areas at the center which explains their position and genomic activity. Regions important for chromatin conformation, such as TAD boundaries and nucleosomal regions show a significant variation in presence of motifs found by DeepBend, which hints that these bendability motifs play a role in the higher order organization of chromosome as well.

Such a simple but effective interpretable model provides the scope to easily visualise patterns’ underlying properties in small sequences. We think the design choices made in DeepBend can be instrumental for other sequence-based problems as well, providing us with further understanding of how deep learning models make accurate predictions. Moreover, our study has also put forward the motif patterns and their relative arrangements that are significant for bendability. These can be further studied to understand how sequence features encode the overall shape and geometry of DNA.

## Methods

### 1 Bendability Datasets

Basu et al. ^4^ developed ‘loop-seq’ assay to measure looping rate of short DNA sequences. They defined the term intrinsic cyclizability which was proved to be highly correlated with DNA bendability. We used their measured values of intrinsic cyclizability as a measure of DNA bendability. These values were obtained from the five sequence libraries, which are Cerevisiae Nucleosomal Library, Random Library, Tiling Library, ChrV Library and Library L, provided as supplementary data with their paper.

#### 1.1 Preparation of test datasets from libraries with tiled overlapping sequences

When we trained and tested our model on the same library (see Supplementary Table 3), we avoided overlapping of sequences between training and test dataset. In the ChrV library, adjacent sequences are offset by 7bp. In the Tiling library, adjacent sequences taken from the same 2001 bp region are also offset by 7bp. The overlap between nearby sequences can range from 43bp to 1bp. There is also a correlation between bendability values of adjacent sequences. In ChrV library, adjacent sequences have a Pearson’s correlation (r) of 0.467, 0.329, 0.220, 0.104 and so on. For Tiling library, adjacent sequences have a Pearson’s correlation (r) of 0.666, 0.474, 0.318, 0.159 and so on. Training and testing are done carefully when these libraries are used as datasets. If a sequence is taken into the test set, 7 sequences upstream and downstream and the sequence itself is not taken into the training set. This ensures that there is no leaking of information from the test set into the training set. For the test set, we have taken 4% of the total sequences. The ChrV test set consists of 3,297 sequences from a total of 82,405 sequences. The ChrV training set consists of 44,604 sequences. The rest are lost for avoiding leakage. The Tiling library, its test and train set consists of 82,369, 3,205 and 44,657 sequences respectively.

### 2 DeepBend Model Architecture and Rationale

The primary goal of using a deep learning model is to improve prediction accuracy and improve interpretability. With our model, we were able to separate out small motifs and the patterns of distribution of these motifs that correspond to bendability positively and negatively. The same motifs, depending on how they are arranged in the sequence, can contribute to positive or negative bendability, our model provides more insight into these motif arrangements.

Our model is a 3-layered Convolutional Neural Network (CNN) regression model that predicts the bendability (intrinsic cyclizibility) of a 50bp sequence. One-hot encoded forward and reverse-complement DNA sequences each of length 50bp and width 4 is given as input to the model. The first two layers are shared between the forward and reverse-complement input sequences so that both strands are considered by the model without compromising training time. The element-wise maximum of the outputs from these sequences are then passed to the next layers.

#### Layer 1

The first layer is the custom 1D convolution from the MuSeAM model ^10^ that learns the multinomial distribution of motif patterns. In this multinomial convolutional layer, 256 filters of size 8×4 have been used with “same” padding and stride 1. The filters are applied separately on one-hot encoded forms of both forward and reverse complement sequences. ReLU activation function is applied after convolution operation. The purpose of this layer is to capture small motif patterns. The advantage of using a multinomial convolutional layer is that it provides salient motif patterns without requiring additional post-processing, Methods 3.2. The outputs of the 1st multinomial convolution layer are the positive matching scores of the motifs at each position. Using specificity factors, alpha = 75.0 and beta = 1/75.0, provides confident looking motifs without degrading the accuracy of the model.

#### Layer 2

The 2nd layer is a 1D convolution layer with 2 to 16 filters, each of length 40 and width 256 and applying “same” padding, stride 1. This layer is used to capture patterns of greater lengths. It is L2 regularised (λ=0.0005). The second convolution convolves over the matching scores of all the motifs from the previous layer in segments of the sequence and produces output depending on the relative spatial positioning of these motifs. These convolutions are of length 40 and so they can capture patterns of motif distribution in larger areas. By hyperparameter tuning we have found for the bendability datasets, increasing the length of filters beyond the input sequence length or using more than 16 filters does not increase performance any further. Furthermore, adding padding allows these convolutions to detect the patterns at any position across the sequence, even near the edges, without having to find multiple shifted versions of the same pattern. Each filter of the second layer *F_i_* is a two-dimensional matrix of size *n*_1_ × *l*_1_, where *n*_1_ is the number of filters in the first layer and is the distance over which the patterns are assumed to be periodic. The *j*-th row of *F_i_* corresponds to the *j*-th filter in the first layer and learns the *i*-th first-order relative spatial pattern of the *j*-th motif relevant to bendability. The weights of the filter *F_i_* at each position represent the relative arrangements of the motifs. Each filter *F_i_* represents higher-order distributive patterns of all the motifs together. The model captures the most prominent patterns that strongly influence model performance. Each output feature map of this layer is a vector of length 50 in which a value at a position denotes how well the region around that position in the input sequence matches the arrangements of motifs represented by the corresponding filter.

Element-wise maximum score is taken between the feature maps of this layer generated from forward and reverse-complement input sequences. After that, the ReLU operation is performed. This propagates the best matching scores to the next layers and allows the model to learn to match motifs at a DNA sequence position irrespective of the strand

#### Layer 3

The third layer is a 1D convolution layer with a single filter of width 50 that uses the matching scores with each distribution/arrangement pattern from the previous layer and provides a single floating-point output.

The output of the penultimate layer (second layer in our case) is a 2D matrix *U* of size *n*_2_ × *l*_2_ where *n*_2_ is the number of filters in the second layer and *l*_2_ is the length of the input sequence. Because of using padding throughout the convolutional layers, the lengths of the inputs and outputs are kept the same. We use *U_i,j_* to denote the output for the filter *i*. and position *j*. The final layer is a convolutional layer with a single filter with no padding and with no activation function at the end. So, the final convolution layer does only one convolutional operation to produce the final floating-point output *Y* and does not move across the input matrix on any axis. The input dimensions of the last layer weights *W* correspond to the output dimensions of the previous layer, i.e., *n*_2_ × *l*_2_. The weight *W_i,j_* corresponds to or is applied to the output *U_i,j_*. Our target is to interpret what influences (i.e., increases or decreases) the output *Y*, i.e., what makes the sequence more bendable (output more positive) or more rigid (output more negative). The outputs for each higher-order feature and input position from the previous layer, *U_i,j_*, are non-negative numbers corresponding to how well the feature of filter *i*. matched with the input at the corresponding position *j*. The sign of the weight *W_i,j_* of the last layer determines whether the output *U_i,j_* contributes positively or negatively to the final output, *Y* and its value or magnitude determines the contribution factor. If we look at it in another way, for each input position *j*, every filter from the second layer, *F_i_* has a weight *W_i,j_* assigned to it in the last layer. An output/match for the filter *F_i_* at a position *j* contributes to the final output *Y* by being factored with that weight *W_i,j_*. For example, suppose the weight for filter #12 for input position 5, *W*_12,5_= 0.2. In that case, we can assume that if there is a match of the filter #12 at position 5 then the final output will be moved to the positive direction in proportion to the weight, *W*_12,5_ and the matching output. In the absence of careful regularization, a filter’s contribution may fluctuate between positive and negative values (also of varying magnitudes) across input positions. To limit this, we have added variance regularization ^26^ along the positional axis of the last layer convolution. As a result, the weights for a particular filter would be of similar value and a filter would contribute similarly to the final score if it matches any of the positions. Thus we can clearly say in which direction a match of the filter pattern *F_i_*, i.e, the filter itself is contributing, from the corresponding weights in the last layer, 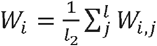. Then, we can identify the distributive patterns of motifs that contribute positively and negatively to bendability and also their relative magnitude of contribution. The last layer is a convolutional layer instead of a fully connected layer so that we do not lose the positional and filter dimensions. Variance regularisation loss for final layer weights *W* is calculated as follows:

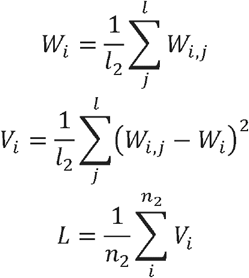

Since all the patterns are in the filters of a single layer, we can get the higher-order relations by observing filter weights. We do not need to do a post-hoc analysis ^27^ on our model for finding out the variation of output due to the relative arrangement of motifs.

### 3 Extracting motif pattern from trained model

#### 3.1 Multinomial Convolution Operation

The convolution layer in MuSeAM learns multinomial distributions. For this, the convolution matrix *W* is transformed into a multinomial distribution matrix *T* where:

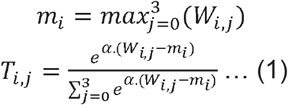

and *α* is a free parameter that determines the specificity of the motif. Here *i* and *j* are indices for the rows and the columns of *T*, respectively. For each multinomial convolution matrix of size *T* and a one-hot encoded nucleotide sequence of length *L*, a convolution operation computes the term 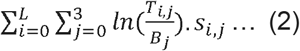, on each *L*-length subsequence *s* of *S*. where *B* is a background distribution over the four DNA nucleotides. The log-likelihood vector is then passed through a ReLU operation which only passes those values which pass a certain threshold, i.e those that have a good enough match. The second convolution layer helps capture the distribution of the motifs in the sequence. The output of the 1st multinomial convolution layer is the matching score of the motifs at each position.

#### 3.2 Extracting motif patterns from multinomial filters

For getting the motif patterns from the convolution matrix, transformation (1) is used to get the probability matrix for the motifs.

### 4 Ranking motif by global importance analysis

In order to understand the importance of each of the motifs obtained from our model towards bendability, we check the average change in predicted bendability score for sequences in a dataset when the motifs signals are turned off ^28^. The first convolutional layer and the ReLU gives the matching scores of each motif. By turning the outputs of one of the motifs to zeros, the model provides output as if the motif were not present in the sequence. If *X* are the input sequences in the dataset *f*: *x* → *y* is the function of the original model and *f_motif_*: *x* → *y* is the function of the model with the output of a motif turned off then the importance of that motif is, 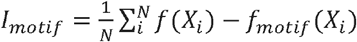. We used the sequences from the ChrV Library for determining the global importance of motifs (Supplementary Material 2).

### 5 Motif strong/weak matching score

The matching scores of motifs are obtained from the output of the first layer of our model. The output of the first layer is the log-likelihood of the sequence segment being the motif sequence with respect to some background distribution. If the log-likelihood crosses a certain threshold we say that there is a match. That threshold is applied by the ReLU operation after the convolution. We call this output the matching score of a motif, which is obtained at each position for all input sequences. For finding the matching scores of a chromosome of S. Cerevisiae we used the 50bp sequences from that chromosome offset at 7bp from the adjacent ones. We ran the model on these sequences and obtained the output from the 1st layer, which are matching scores of the motifs of each 50bp sequence at each position. Then we applied ReLU activation function and averaged the overlapping scores to get the matching score of the entire chromosome.

### 6 Denoting Bendability across chromosome

We derived the mean bendability across chromosome V as follows: ChrV library contains bendability values of 50-bp sequences at 7 bp offset. We calculated bendability value at each bp by taking the average of the overlapping 50-bp sequences passing that location. When predicting cyclizability with DeepBend in a chromosome, we similarly used 50-bp DNA sequences at 7 bp offset as input to our model.

### 7 Boundary and Nucleosome Position Determination

We obtained yeast chromatin interaction data found with Micro-C XL ^29^. From this data, we determined domains and domain boundaries at 200-bp resolution with Hi-C Explorer (calling options: min_depth = 1000, max_depth = 10000, step = 1000, thres_comparison = 0.05, delta = 0.01, correct_for_multiple_testing = fdr). In 16 chromosomes of yeast, we found 2862 200-bp long boundaries in total (Supplementary Table 4).

We obtained nucleosome dyad locations from ^30^. A nucleosome was considered as the region from −73 bp to +73 bp from the dyad, totalling 147 bp. The rest of the DNA sequence was considered as linkers. We downloaded nucleotide sequences of all chromosomes of Yeast from the Saccharomyces Genome Database (SGD) ^31^. We also downloaded Yeast gene transcripted regions from YeastMine of SGD. We considered the whole transcripted regions as gene.

### 8 Comparing Bendability of Boundaries and Domains

With Hi-C Explorer, we determined 200-bp long boundaries in all 16 chromosomes of Yeast (Methods 7) and denoted the rest of the chromosome as domain regions. So, in a chromosome with N boundaries, we had N+1 domains. For each boundary, we determined its middle bp and took −499 bp to +500 bp from this middle bp to check bendability of boundary regions. To compare bendability of domains and boundaries, we also took 1000 bp sections in domains. When the length of a domain, L, was not a multiple of 1000, we excluded (L mod 1000) / 2 bp from each flank of domain and sectioned the rest. We then took bendability value of boundaries and domain sections according to Method 6.

### 9 Bendability Quotient

The bendability quotient of a k-mer in a dataset is defined as the average bendability of all the sequences from a dataset that contains the k-mer ^5^. We have calculated the bendability quotient using the Random Library as number of k-mers should theoretically be most eqaually distributed here.

## Supporting information

All Supplemental Materials

## Code and Data Availability

The required data were obtained from ^4^ and the code will be released upon the manuscript’s acceptance.

## Supplementary Notes

### 1 Different Models we have tried

#### 1.1 MuSeAM model

MuSeAM model detects the presence of motifs in the sequence. But fails to capture how these motifs are spread in the sequence. Training with larger motifs in a single layer means that they have to capture combinations of motifs and their distributions. This makes the model larger and harder to interpret. Models we have trained in this architecture have not reached the expected results. (r: 0.64). We have also tried using filters of different lengths in our MuSeAM model in order to allow capturing larger regions. (r: 0.781).

#### 1.2 Model with dinucleotide encoded input

To show that the distribution of only dinucleotides is not enough to accurately predict bendability we also trained a CNN model that takes dinucleotide one-hot encoding as input and learns dinucleotide occurrence patterns in the first 1D convolution layer and their distribution in the second convolution layer. This model achieves Pearson’s r=0.802 on the test dataset whereas a previous model which takes dinucleotide counts and gapped dinucleotides counts achieved Pearson’s r=0.6.

#### 1.3 Multinomial CNN-RNN Model

We have also tried increasing and decreasing model depth, width and changing hyperparameters. We had also experimented with RNNs in our second layer. Using a model with multinomial CNN layer and then a bidirectional GRU layer resulted in Pearson’s correlation (r) of 0.8452 in the test set. Although the models are very close in their performance, we had decided to move forward with the DeepBend architecture as it provided the scope for a better understanding of what makes sequences more or less bendable. All the models have been trained on Tiling library, validated on ChrV test library (Methods 1) and tested on Random library. The results of these models are summarised in Supplementary Table 2.

#### 1.4 Machine Learning Models using sequence features

We used several machine learning models such as Linear Regression, Support Vector Machine and Random Forest to predict bendability of sequences from extracted features. As features, we used the number of times each of the 16 dinucleotides, 64 trinucleotides and 256 tetranucleotides occurred in a sequence and 136 helical separation extent values, which is a measure of tendency of a dinucleotide pair to be separated at helical distance ^16^. Thus, each sequence was converted into a 152 feature vecto. The models were trained on Tiling library and tested on Random library, Cerevisae nucleosomal library and chromosome V library. The Pearson’s correlation between predicted and actual values for these models are shown in Supplementary Table 1.

### 2 Motifs in Chromatin Conformation Found by Meme-suite

We used streme [15] to find enriched motifs of 8-bp width in domains and boundaries. −250 bp to +250 bp around all boundaries middle in 16 chromosomes were considered as boundary sequences. The rest of the chromosome was considered in domains. Average domain sequence length was about 3700bp. When finding enrichment of boundary sequences, we used domain sequences as control sequences and vice versa.

**Supplementary Fig. 1.**
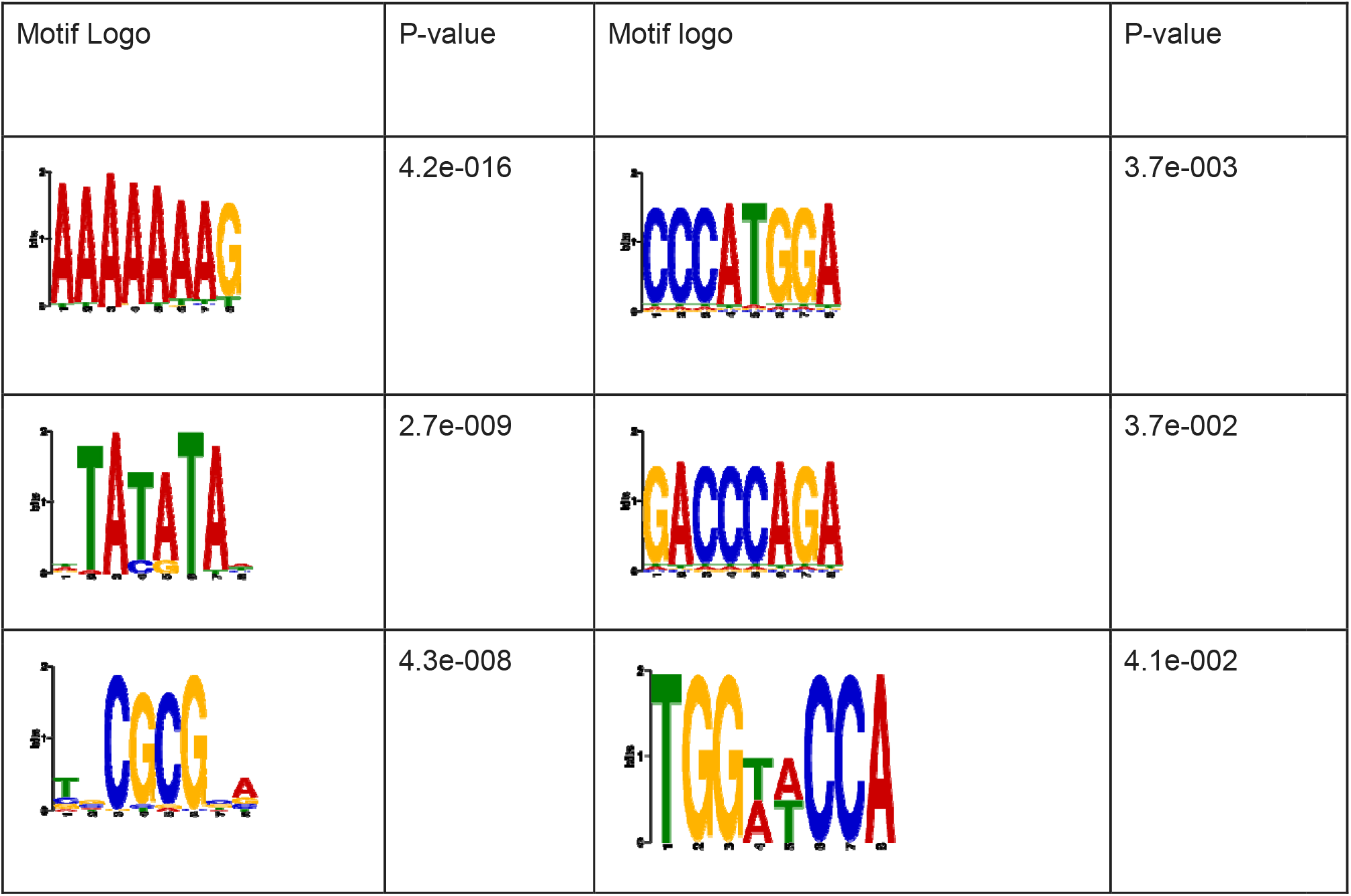
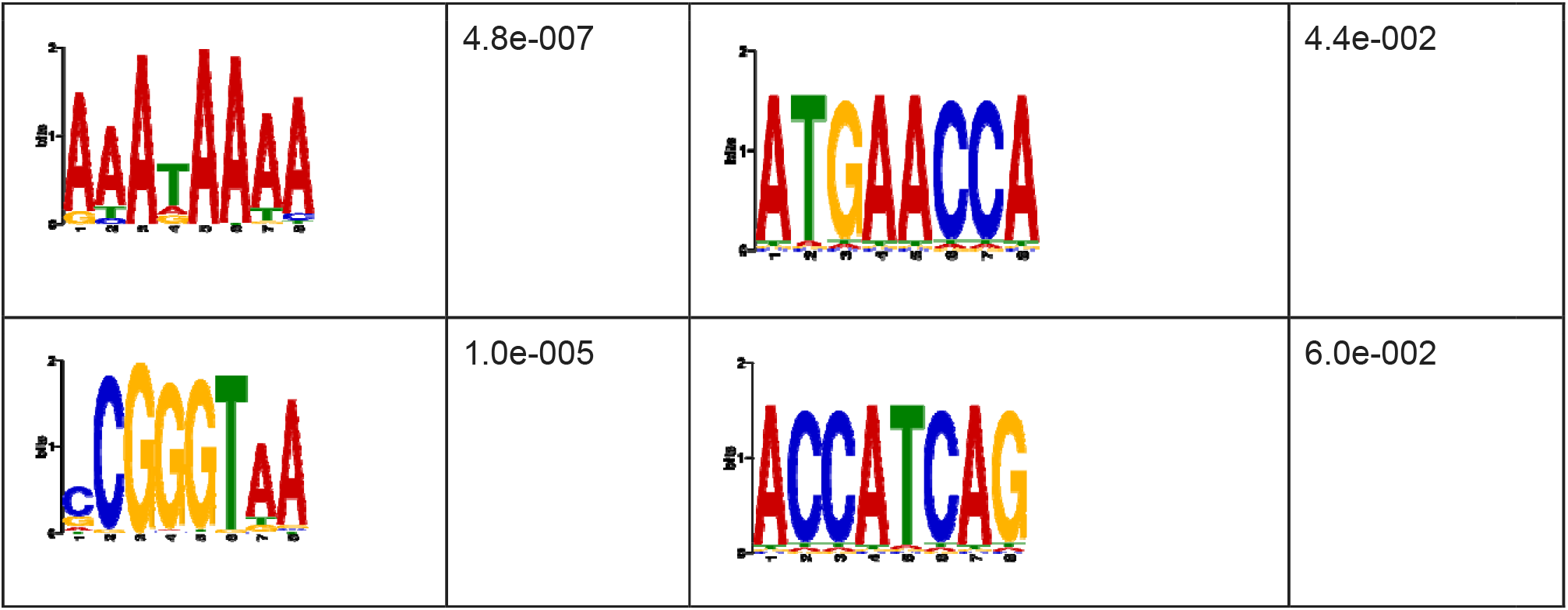
Motifs found by meme-suite. (Left) Motifs that are relatively enriched in boundaries compared to domains. (Right) Motifs that are relatively enriched in domains compared to boundaries.

