## Supplementary material for "DeepBend: An Interpretable Model of DNA Bendability": All Supplemental Materials: List of Supplementary Materials.docx

###### **Materials 1:**

Figures of 256 motifs.

###### **Materials 2:**

Ranking of motifs by Global Importance Analysis.

###### **Materials 3:**

Z-values of motif matching score distributions in boundaries and domains.

###### **Materials 4:**

A zipped archiver of our codes and the data necessary to reproduce our results. The zipped archive also contains a README file describing the dependencies and commands to run our codes.
