## Supplementary material for "DeepBend: An Interpretable Model of DNA Bendability": All Supplemental Materials: Supplementary Tables.docx

###### **Supplementary Table 1: Pearson’s correlation (r) of true and predicted values from different models when tested on different datasets. The models have the trained on Tiling dataset.**

| Model Name | Performance on Training dataset | Performance on Test dataset | | |
| --- | --- | --- | --- | --- |
|  | Tiling library | Random library | Cerevisiae Nucleosomal library | ChrV library |
| Ridge Regression | 0.56 | 0.4 | 0.46 | 0.38 |
| SVM | 0.55 | 0.41 | 0.46 | 0.38 |
| Random Forest | 0.84 | 0.29 | 0.33 | 0.25 |
| XGBoost | 0.99 | 0.2 | 0.25 | 0.19 |
| Neural Network Model with Feature inputs | 0.62 | 0.41 | 0.48 | 0.4 |

###### **Supplementary Table 2: Pearson’s correlation (r) of true and predicted values from DeepBend model trained on Tiling and tested on Random library**

| Model | Train | Test |
| --- | --- | --- |
| MuSeAM | 0.7694 | 0.6401 |
| MuSeAM with multi-sized filters | 0.9085 | 0.781 |
| Dinucleotide encoded model | 0.929 | 0.8024 |
| Multinomial CNN + Bidirectional GRU | 0.9586 | 0.8452 |
| DeepBend | 0.94 | 0.8946 |

###### **Supplementary Table 3: Pearson’s correlation (r) of true and predicted values from DeepBend model trained and tested on different datasets**

| trained on | tested on | train | test | whole ChrV | whole random | whole nucleosomal | whole tiling |
| --- | --- | --- | --- | --- | --- | --- | --- |
| ChrV train set | ChrV test set (avoiding overlaps) | 0.755 | **0.74** | **0.753** | 0.85 | 0.878 | 0.87 |
| Random train set | Random test set | 0.905 | **0.841** | 0.733 | **0.902** | 0.88 | 0.86 |
| Nucleosomal train set | Nucleosomal test set | 0.92 | **0.883** | 0.699 | 0.816 | **0.918** | 0.822 |
| Tiling train set | Tiling test set | 0.901 | **0.889** | 0.749 | 0.862 | 0.9 | **0.9** |

###### **Supplementary Table 4: Yeast Chromosome Data**

| Chromosome | Total length (bp)* | Number of boundaries |
| --- | --- | --- |
| I | 230217 | 48 |
| II | 813184 | 187 |
| III | 316618 | 73 |
| IV | 1531930 | 371 |
| V | 576871 | 134 |
| VI | 270159 | 59 |
| VII | 1090937 | 264 |
| VIII | 562640 | 133 |
| IX | 439888 | 104 |
| X | 745746 | 178 |
| XI | 666814 | 166 |
| XII | 1078176 | 226 |
| XIII | 924428 | 234 |
| XIV | 784330 | 194 |
| XV | 1091287 | 259 |
| XVI | 948060 | 232 |
|  |  | 2862 |
| * Total length was less than the original length as we measured the bendability of the whole chromosome using 50-bp segments at 7-bp offset. | | |
